# Critical analysis of radical scavenging properties of atorvastatin in methanol recently estimated via density functional theory

**DOI:** 10.1101/2022.06.28.497928

**Authors:** Ioan Bâldea

**Author notes:** Corresponding author; *Email address* (Ioan Baldea).

## Abstract

In this communication we draw attention on serious flaws that plague recently reported antioxidant properties of atorvastatin (ATV) in methanol. First and foremost, we emphasize that the O-H bond dissociation energies (BDE) of about 400kcal/mol previously reported are completely wrong. Further, we present results refuting the previous claim that the proton affinity (PA) of ATV is smaller than that of the ascorbic acid. That unfounded claim relies on incorrect data for PA’s ascorbic acid (which we correct here) circulated in the literature. Further, we correct the values of the chemical reactivity indices (e.g., chemical hardness, electrophilicity index, electroaccepting and electrodonating powers), which were inadequately estimated previously via Kohn-Sham HOMO and LUMO energies. Finally, our updated values for O–H bond dissociation enthalpy (BDE = 91.4 kcal/mol) and electron transfer enthalpy (ETE = 105.7kcal/mol) tentatively suggest that direct H-atom transfer (HAT) and sequential proton loss electron transfer (SPLET) may coexist.

## 1. Introduction

Atorvastatin (ATV) [1] is one of the best selling drug belonging to the class of statins [2, 3] widely administered over more than 25 years in a variety of settings to lower the “bad” cholesterol and fats —e.g., low-density lipoprotein (LDL), triglycerides —, decrease the risk of heart disease, prevent strokes and heart attacks, etc [4, 5, 6, 7].

Quantum chemistry can provide important insight into the antioxidant activity of ATV, which still remained elusive. For this reason, the first efforts in this direction undertaken only recently [8] are certainly welcome. Unfortunately, the results presented in ref. [8] are plagued by serious flaws, and drawing attention to this fact is an important aim of the present communication. As brief justification of this assertion suffice it to mention what is undoubtedly the most eye-catching error of ref. [8]. For ATV’s O–H and N–H bonds, values of about 400kcal/mol (> 17 eV) are reported (cf. Table 2 of ref. [8]). Roughly, these are four times larger than any BDE value characterizing the well documented O–H and N–H groups [5], which represent mandatory ingredients of any radical scavenger just because of their well known low BDE’s. The aforementioned values even by far exceed the strongest chemical bonds known today, amounting to (generously speaking) ~ 10 eV.

**Table 1:** Electronic energy *E*_0_, zero point vibrational energy *ZPE*, and thermal corrections to enthalpy *TCH* for neutral ATV and related species in methanol at the B3LYP/6-31+G(d,p) level of theory. For open shell systems, calculations were done within the unrestricted (UB3LYP) framework, and the values of the total spin before (label *b*) and after (label *a*) annihilation of the first spin contaminant are indicated. ATV*^•+^ = ATV cation; ATVwoH^•^ = ATV radical after H-atom removal from 1-OH position (cf. Figure 1).

| Species | $E_0$ | $ZPE$ | $TCH$ | $\langle S^2 \rangle_b$ | $\langle S^2 \rangle_a$ |
| --- | --- | --- | --- | --- | --- |
| ATV | -1864.24567013 | 0.623552 | 0.663158 |  |  |
| ATV$\bullet^+$ | -1864.04563713 | 0.624017 | 0.663523 | 0.7633 | 0.7501 |
| ATVwoH$\bullet$ | -1863.59072281 | 0.610822 | 0.649807 | 0.7633 | 0.7501 |
| ATVwoH$^-$ | -1863.78861771 | 0.610362 | 0.649465 | | |

**Table 2:**
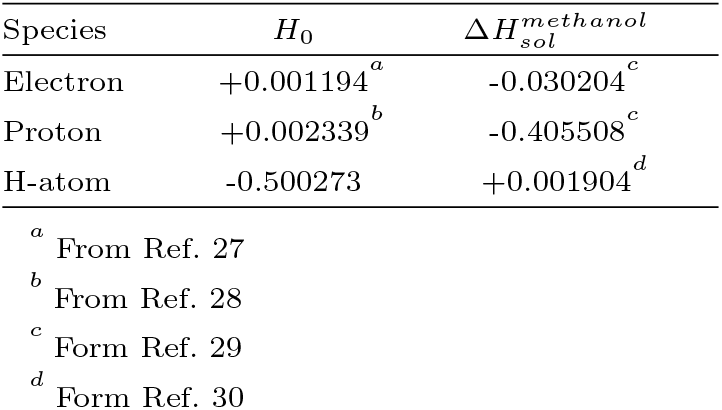
Gas phase enthalpies Ho and solvation enthalpies Δ*H_sol_* in hartree needed in the present calculations. Except for the gas phase enthalpy of the hydrogen atom computed by us using B3LYP/6-31+G(d,p), all other entries are literature data [30, 27, 28, 29].

## 2. Computational details

The quantum chemical calculations based on the density functional theory (DFT) done in conjunction with this study employed the GAUSSIAN 16 [9] suite of programs on the bwHPC platform [10]. We used the three parameter B3LYP hybrid DFT/HF exchange correlation functional [11, 12, 13, 14] along with the Pople 6-31+G(d,p) basis sets [15, 16]. Solvent (specifically, methanol) effects were accounted for within the polarized continuum model (PCM) [17] using the integral equation formalism (IEF) [18].

For open shell radicals (ATV^•,+^ and ATVwoH^•^, cf. Tables 1, S2, and S3) we performed unrestricted (UB3LYP) calculations. Therefore, it is worth mentioning that, similar to other similar cases [19, 20, 21, 22], we can safely rule out spin contamination artifacts (cf. Table 1).

Because literature studies on antioxidant activity often use optimized geometries and vibrational corrections due to zero point motion in the gas phase with smaller basis sets, and only compute electronic energies with solvent and larger basis sets, we emphasize that all our optimized geometries (presented in Tables S1 to S5) and vibrational corrections due to zero point motion (Table 1) were also computed at the B3LYP/6-31+G(d,p)/IEFPCM level of theory. GABEDIT [23] was used to generate Figures 1 and 2. All thermodynamic properties were calculated at *T* = 298.15 K.

**Figure 1:**
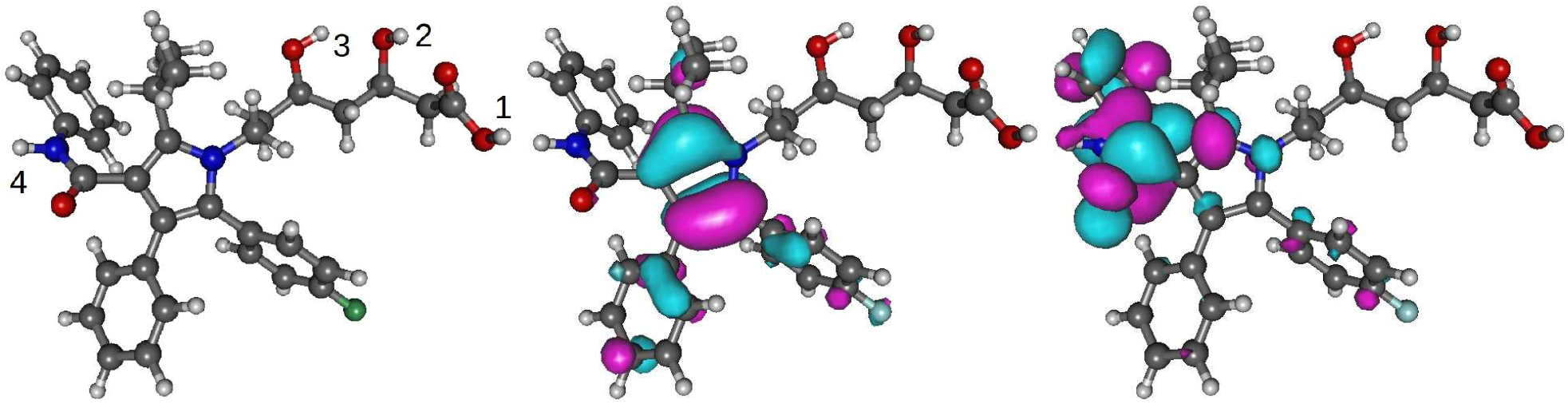
Optimized geometry with labels of the O-H groups (left), HOMO (middle), and LUMO (right) spatial distributions of the atorvastatin molecule (ATV).

**Figure 2:**
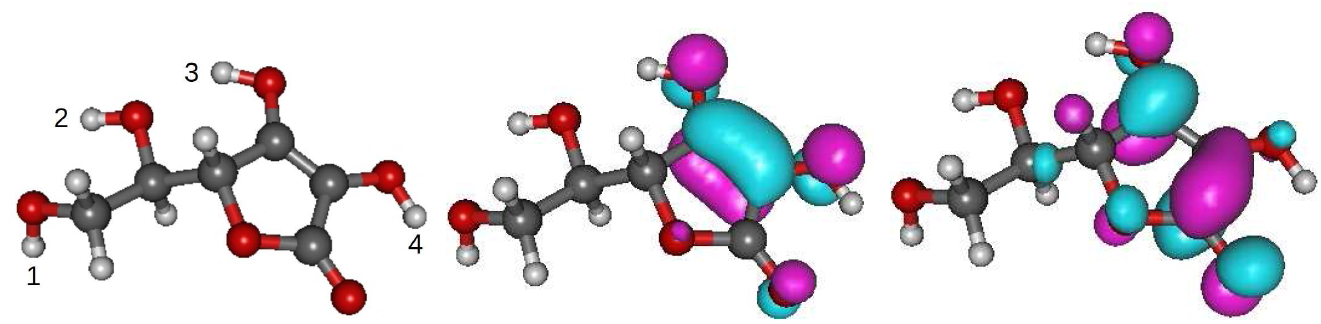
Optimized geometry with labels of the O-H groups (left), HOMO (middle), and LUMO (right) spatial distributions of ascorbic acid.

## 4. Results and discussion

Antioxidants (XH) are molecules that inhibit oxidation processes by transferring a hydrogen atom to free radicals (R). The three main antioxidative mechanisms (HAT, SET-PT, and SPLET) along with the pertaining reaction enthalpies (BDE, IP and PDE, PA and ETE, respectively) are depicted below.

Direct hydrogen atom transfer (HAT):

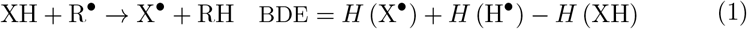

Stepwise electron transfer proton transfer (SET-PT):

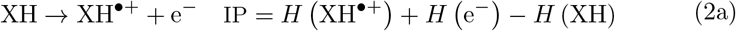

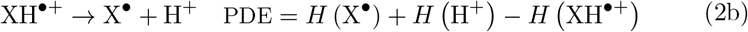

Sequential proton loss electron transfer (SPLET):

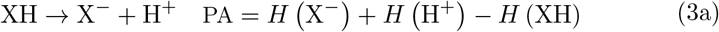

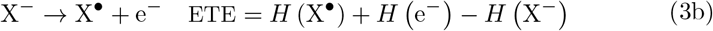

The thermodynamic parameters listed above (BDE, IP and PDE, PA and ETE) provide useful information needed to assess the radical scavenging activity of ATV. They are enthalpies of reaction and can be obtained from standard Δ-DFT quantum chemical calculations [24, 25, 26]).

In this communication, we will restrict ourselves to the O-H bond of the carboxylic acid group (1-OH position in Figure 1), which, out of the three O-H bonds of ATV, has according to ref. [8] the lowest PA and ETE. In the context of ref. [8], the lowest ATV’s PA value is not only relevant for the SPLET mechanism (claimed to be the most efficient pathway in ATV [8]) but also because of the claim [8] that it is lower than that of ascorbic acid (vitamin C). To refute this claim, we also re-estimated PA for ascorbic acid and emphasize the difference from an inadequate value circulated in the literature. Unlike that value shown in ref. [8], our PA for ascorbic acid was computed at the same B3LYP/6-31+G(d,p)/IEFPCM level of theory also employed for ATV.

The presently calculated enthalpies of the ATV species entering the righthand side of eqs. (1), (2), and (3) are included in Table 1. To estimate the various reaction energies, the gas phase enthalpies of the hydrogen atom, proton, and electron as well as their solvation enthalpies are also needed. They are listed in Table 2.

Insertion of the values of Tables 2 and 1 into eqs. (1), (2), and (3) yields the desired ATV’s thermodynamic parameters. They are given in Table 3 and visualized in Figure 3 along with the previous values (written in italics) of ref. [8]. As visible there, in contrast to the huge value (403.8kcal/mol), our value for ATV’s O-H BDE (91.4 kcal/mol) has nothing extravagant; it lies in the range typical for hydroxy groups [5].

**Figure 3:**
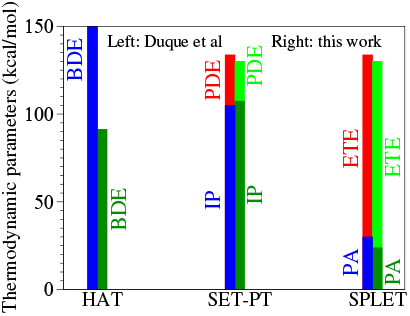
Thermodynamic parameters quantifying the radical scavenging activity of ATV in methanol as estimated in the present study along with those reported in ref. [8]. The underlying numerical values were taken from Table 3.

**Table 3:** O–H bond dissociation enthalpies (BDE), ionization enthalpies (IP), proton dissocia-tion enthalpies (PDE), proton affinity (PA), and electron transfer enthalpies (ETE) computed for ATV and ascorbic acid in methanol at the B3LYP/6-31+G(d,p) level of theory. Values previously reported in ref. [8] are shown in italics. All quantities in kcal/mol.

|  | HAT | SET-PT |  | SPLET |  |
| --- | --- | --- | --- | --- | --- |
|  | BDE | IP | PDE | PA | ETE |
| Present work | 91.4 | 107.0 | 22.4 | 23.8 | 105.7 |
| Ref. [8] | *403.8* | *105.1* | *28.8* | *30.1* | *103.8* |

Although neither the finding on incorrect BDE value previously reported [8] nor our updated BDE value should be too surprising for scholars familiar with O-H groups, the aforementioned is also important because it challenges the claim of ref. [8] that SPLET is the dominant mechanism responsible for the radical scavenging activity of ATV. This claim on the SPLET prevalence is hardly substantiated by the updated value BDE= 91.4 kcal/mol, “comfortably” lower than ETE=106.2 kcal/mol (second step of SPLET, cf. eq. (3b)) and also lower than IP=107.0 kcal/mol (first step of SET-PT, cf. eq. (2a)). In view of our results (Table 3 and Figure 3), we suggest that HAT rather than SPLET is the ATV’s dominant antioxidant mechanism in methanol.

Our proton affinity (PA=23.8 kcal/mol, cf. Table 3) is even lower than the previous estimate (PA=30.1 kcal/mol [8]). Nevertheless, as elaborated below, we refute the claim [8] that this value is smaller than for ascorbic acid.

To support that claim, ref. [8] quoted a value PA=34.2 kcal/mol for ascorbic acid taken from literature but did not mention to which O-H group this value belongs. This is a significant issue because there are four O-H groups in ascorbic acid (cf. Figure 2). To elucidate this point, we also performed B3LYP/6-31+G(d,p)/IEFPCM calculations for all the four O-H groups of ascorbic acid in methanol. Our results (to be reported in detail elswhere) settles the confusion. Up to minor difference understandable in view of the different levels of theory, the value of PA=34.2 kcal/mol for ascorbic acid presented in ref. [8] corresponds to the 4-OH position (cf. Figure 2), for which we estimated PA=35.0kcal/mol. However, this is not the lowest PA of ascorbic acid. The lowest PA of ascorbic acid determined by us (20.5 kcal/mol) corresponds to the 3-OH position.

A detailed comparison between our data for ascorbic acid and those from previous literature including ref. [8] deserve separate consnideration. What matters from the present discussion is that the lowest PA value (20.5 kcal/mol) of ascorbic acid is lower than the ATV’s lowest value (PA=23.8 kcal/mol) presently considered.

Ref. [8] also reported global chemical reactivity indices for ATV in methanol: chemical hardness *η* ≡ *E_g_*/2, chemical softness *S* ≡ 1/*E_g_*, electronegativity *χ* ≡ (IP + EA)/2, electrophilicity index *ω* ≡ *χ*^2^/(2η) as well as electroaccepting *ω*^+^ ≡ (IP + 3EA)^2^/(16Eg) and electrodonating *ω*^-^ ≡ (3IP + EA)^2^/(16Eg) powers. Here, *E_g_* ≡ IP-EA is the fundamental (or transport) “HOMO-LUMO” gap [31, 32, 26]. What makes their estimates of Ref. [8] highly questionable is the determination of the ionization energy (IP) and electroaffinity (EA) from the Kohn-Sham (KS) HOMO and LUMO energies obtained from DFT calculations in solvent 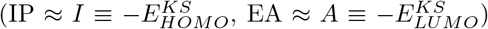. It is well known that the KS gap *I* — *A* differs from *E_g_* even if computed with the exact exchange-correlation functional [33, 32], and the presence of a solvent adds a further difficulty [34].

To estimate the global chemical reactivity indices, we computed the electroaffiity EA = *H* (XH) + *H* (e^−^) — *H* (XH^•−^) from the enthalpy of the anion (XII^•−^ ≡ ATV^•−^), i.e., as counterpart of of IP of eq. (2a). As expected, our values collected in Table 4 substantially differ from those previously reported [8].

**Table 4:** Presently computed global chemical reactivity indices (in eV) substantially differ from those estimated in ref. [8] shown in italics.

| | IP | EA | $E_g$ | $\chi \equiv -\mu$ | $\eta$ | $S$ | $\omega$ | $\omega^+$ | $\omega^-$ |
| --- | --- | --- | --- | --- | --- | --- | --- | --- | --- |
| Present work | 4.64 | 0.72 | 3.92 | 1.96 | -2.68 | 0.26 | 1.83 | 0.74 | 3.42 |
| Ref. [8] | *5.75* | *1.04* | *4.71* | *2.35* | *-3.40* | *0.21* | *2.45* | *1.05* | *4.44* |

## Conclusion

In closing, clearly contradicting the main conclusions of ref. [8], the results presented above demonstrated that the O-H bonds of atorvastatin (ATV) are by no means unusually strong, and that the lowest proton affinity of ascorbic acid is lower than the lowest proton affinity of ATV. In addition, we showed that the global (adiabatic) chemical reactivity indices based on IP and EA values reliably determined via enthalpies of reactions substantially differ from those based on the Kohn-Sham HOMO and LUMO energies.

Last but not least, the presently corrected value of the O-H bond dissociation energy, smaller than the second step of the SPLET (BDE = 91.4kcal/mol versus ETE = 105.7 kcal/mol), may tentatively suggest a bottleneck effect played by the second step of the SPLET pathway and herewith a nontrivial interplay with direct hydrogen atom transfer (HAT) in determining the antioxidant activity of ATV in methanol.

## Acknowledgments

The author thanks Ederley Vélez Ortiz for the kindness in providing him valuable details related to her recent work [8]. Financial support from the German Research Foundation (DFG Grant No. BA 1799/3-2) in the initial stage of this work and computational support by the state of Baden-Württemberg through bwHPC and the German Research Foundation through Grant No. INST 40/575-1 FUGG (bwUniCluster 2.0, bwForCluster/MLS&WISO 2.0, and JUSTUS 2.0 cluster) are gratefully acknowledged.

## Appendix

**Table S1:** Coordinates of ATV optimized in methanol via RB3LYP/6-31+G(d,p)/IEFPCM. All values in angstrom.

| Atom | X | Y | Z | Atom | X | Y | Z |
| --- | --- | --- | --- | --- | --- | --- | --- |
| F | -3.10161200 | 4.92704700 | -1.32781200 | C | -0.32790500 | 2.57709400 | -1.35193100 |
| O | -6.57690300 | -1.23310100 | 1.55261700 | H | 0.44269000 | 2.22333700 | -2.02930000 |
| O | 4.41731700 | 0.32594500 | 2.25091400 | C | -1.22961800 | 3.54944900 | -1.79029700 |
| N | 0.25030100 | -0.25509300 | 0.75806400 | H | -1.17733000 | 3.96041800 | -2.79273000 |
| C | -6.82160200 | -0.44128000 | 0.65138800 | C | -2.32369000 | 3.48862800 | 0.38698700 |
| C | 3.86938900 | -0.35311100 | 1.36852800 | H | -3.09917400 | 3.86083200 | 1.04770900 |
| C | 1.39000000 | -0.96319700 | 1.09384600 | C | -1.40634700 | 2.52375600 | 0.80931800 |
| C | 0.59359200 | 1.04694000 | 0.40288400 | H | -1.46479800 | 2.15264100 | 1.82800500 |
| C | -2.21114900 | 3.97957500 | -0.90732300 | C | -1.10282800 | -0.80924800 | 0.60291400 |
| C | 4.33941000 | -2.19429600 | -0.34298900 | H | -1.81198800 | -0.06894900 | 0.97457100 |
| C | 2.46907000 | -0.09340200 | 0.94903000 | H | -1.19074400 | -1.68262200 | 1.24364900 |
| C | 1.96955300 | 1.18500800 | 0.52827000 | C | -3.98389200 | -0.99399800 | -0.65294500 |
| C | -0.40019100 | 2.04821300 | -0.05111300 | H | -3.88834100 | -0.02560700 | -1.15958400 |
| C | 2.76038900 | 2.41096900 | 0.27950500 | H | -3.97630900 | -0.80376900 | 0.42705600 |
| C | 0.77009300 | -2.42549900 | 3.08542600 | C | -6.53122200 | -0.68786500 | -0.81020800 |
| H | 1.19347100 | -1.66684000 | 3.75129200 | H | -7.43468400 | -1.14786400 | -1.23125000 |
| H | -0.31348200 | -2.27576900 | 3.05035100 | H | -6.39312900 | 0.26697000 | -1.32380100 |
| H | 0.95182800 | -3.40966800 | 3.53092400 | O | -2.75986800 | -3.10006300 | -0.30287200 |
| C | 4.88844400 | -3.48661300 | -0.34505400 | H | -3.68170000 | -3.41604300 | -0.27105500 |
| H | 5.43269900 | -3.83815300 | 0.52720200 | O | -5.53024400 | -2.90105100 | -0.43216300 |
| C | 4.73504000 | -4.31718800 | -1.45643900 | H | -5.81481400 | -2.72152400 | 0.48344000 |
| H | 5.16739000 | -5.31325600 | -1.44104900 | O | -7.42117700 | 0.73769500 | 0.87605400 |
| C | 4.02318300 | -3.87489700 | -2.57572800 | H | -7.62466500 | 0.80861600 | 1.82669300 |
| H | 3.89861600 | -4.52186900 | -3.43850800 | N | 4.57984700 | -1.38251800 | 0.79596400 |
| C | 3.48057200 | -2.58617600 | -2.57438000 | H | 5.45749000 | -1.55457100 | 1.27532400 |
| H | 2.93677500 | -2.22360200 | -3.44185400 | C | 1.42256300 | -2.35903400 | 1.68547100 |
| C | 3.64189500 | -1.74201900 | -1.47322700 | H | 2.48737100 | -2.55783400 | 1.84145800 |
| H | 3.22789500 | -0.74191300 | -1.49676100 | C | 0.91213100 | -3.48775000 | 0.76412600 |
| C | 2.37032200 | 3.64855500 | 0.82485900 | H | -0.17116100 | -3.46314100 | 0.62282600 |
| H | 1.48228200 | 3.69984500 | 1.44686600 | H | 1.38776000 | -3.43813800 | -0.21976100 |
| C | 3.11586300 | 4.80643400 | 0.58994200 | H | 1.16264100 | -4.45635300 | 1.21043400 |
| H | 2.79422300 | 5.74881400 | 1.02466500 | C | -1.42112600 | -1.18590100 | -0.85320900 |
| C | 4.27557900 | 4.75359100 | -0.19014500 | H | -0.64463300 | -1.86432300 | -1.22262900 |
| H | 4.85736400 | 5.65285600 | -0.37037800 | H | -1.38982800 | -0.29025500 | -1.48271800 |
| C | 4.68062300 | 3.52977600 | -0.73262300 | C | -2.78024500 | -1.86891300 | -1.04375000 |
| H | 5.57865800 | 3.47361900 | -1.34153900 | H | -2.87280900 | -2.10211400 | -2.11710700 |
| C | 3.93034100 | 2.37432200 | -0.50176600 | C | -5.32972000 | -1.61629800 | -1.05953500 |
| H | 4.25330800 | 1.43572900 | -0.94222800 | H | -5.30349800 | -1.82877300 | -2.13351500 |

**Table S2:** Coordinates of ATV^•+^ radical cation optimized in methanol via UB3LYP/6-31+G(d,p)/IEFPCM. All values in angstrom.

| Atom | X | Y | Z | Atom | X | Y | Z |
| --- | --- | --- | --- | --- | --- | --- | --- |
| F | -2.544038 | 5.259388 | -1.397056 | C | 0.131406 | 2.828285 | -1.232676 |
| O | -6.630600 | -1.027423 | 1.570355 | H | 0.997283 | 2.521115 | -1.807518 |
| O | 4.391284 | -0.057308 | 2.298101 | C | -0.663418 | 3.867514 | -1.703098 |
| N | 0.205551 | -0.184885 | 0.782218 | H | -0.431193 | 4.383314 | -2.627703 |
| C | -6.894429 | -0.249615 | 0.662324 | C | -2.123976 | 3.615475 | 0.243017 |
| C | 3.846333 | -0.635431 | 1.352445 | H | -2.983344 | 3.959546 | 0.807057 |
| C | 1.295381 | -0.987486 | 1.106190 | C | -1.339967 | 2.558674 | 0.690408 |
| C | 0.651578 | 1.056695 | 0.417589 | H | -1.581913 | 2.092796 | 1.638661 |
| C | -1.773677 | 4.239793 | -0.951944 | C | -1.185210 | -0.667053 | 0.607185 |
| C | 4.091951 | -2.438099 | -0.424394 | H | -1.856546 | 0.093024 | 0.997898 |
| C | 2.469539 | -0.223356 | 0.919347 | H | -1.307618 | -1.546749 | 1.230729 |
| C | 2.099035 | 1.065834 | 0.523379 | C | -4.061815 | -0.801008 | -0.657196 |
| C | -0.203557 | 2.145058 | -0.040532 | H | -3.971836 | 0.161481 | -1.175716 |
| C | 2.964472 | 2.223721 | 0.306531 | H | -4.048939 | -0.599689 | 0.420817 |
| C | 0.558944 | -2.354523 | 3.101888 | C | -6.611839 | -0.508671 | -0.798711 |
| H | 1.029930 | -1.616543 | 3.757256 | H | -7.515687 | -0.976545 | -1.210030 |
| H | -0.511970 | -2.140876 | 3.057594 | H | -6.481826 | 0.441881 | -1.322326 |
| H | 0.683890 | -3.342842 | 3.553977 | O | -2.834991 | -2.905620 | -0.303086 |
| C | 4.413724 | -3.803653 | -0.417301 | H | -3.757839 | -3.218900 | -0.255863 |
| H | 4.901516 | -4.235927 | 0.451473 | O | -5.595224 | -2.712582 | -0.404943 |
| C | 4.102379 | -4.601597 | -1.518764 | H | -5.877165 | -2.525507 | 0.510141 |
| H | 4.357338 | -5.656723 | -1.502024 | O | -7.510438 | 0.922246 | 0.877661 |
| C | 3.457218 | -4.050133 | -2.630688 | H | -7.710329 | 1.000579 | 1.828503 |
| H | 3.209781 | -4.672771 | -3.484604 | N | 4.471147 | -1.664261 | 0.709696 |
| C | 3.143422 | -2.688601 | -2.637800 | H | 5.337137 | -1.937498 | 1.164242 |
| H | 2.658512 | -2.244633 | -3.501789 | C | 1.225119 | -2.367246 | 1.702936 |
| C | 3.469010 | -1.876913 | -1.547734 | H | 2.270470 | -2.643117 | 1.865697 |
| H | 3.249646 | -0.817250 | -1.586653 | C | 0.632122 | -3.459429 | 0.780953 |
| C | 2.591308 | 3.507522 | 0.766566 | H | -0.444916 | -3.360914 | 0.633897 |
| H | 1.654633 | 3.641974 | 1.295001 | H | 1.120036 | -3.458528 | -0.197264 |
| C | 3.440894 | 4.596965 | 0.588768 | H | 0.816821 | -4.431401 | 1.247972 |
| H | 3.147227 | 5.572466 | 0.962992 | C | -1.498004 | -1.000310 | -0.858986 |
| C | 4.667620 | 4.434076 | -0.063368 | H | -0.727161 | -1.675231 | -1.245171 |
| H | 5.324639 | 5.285997 | -0.208009 | H | -1.467455 | -0.091799 | -1.468952 |
| C | 5.049425 | 3.167865 | -0.526203 | C | -2.860874 | -1.679047 | -1.047550 |
| H | 5.997832 | 3.037244 | -1.037367 | H | -2.950733 | -1.911929 | -2.120551 |
| C | 4.213747 | 2.072520 | -0.334872 | C | -5.408080 | -1.433700 | -1.047524 |
| H | 4.518773 | 1.101287 | -0.708739 | H | -5.388469 | -1.657067 | -2.119338 |

**Table S3:** Coordinates of ATVwoH• radical optimized in methanol via UB3LYP/6-31+G(d,p)/IEFPCM. All values in angstrom.

| Atom | X | Y | Z | Atom | X | Y | Z |
| --- | --- | --- | --- | --- | --- | --- | --- |
| F | -2.394990 | 5.382882 | -1.404039 | C | 0.199618 | 2.866108 | -1.236101 |
| O | -6.730500 | -1.249652 | 1.509079 | H | 1.060689 | 2.535662 | -1.805272 |
| O | 4.324321 | -0.170699 | 2.325261 | C | -0.557101 | 3.934121 | -1.704682 |
| N | 0.157056 | -0.163816 | 0.756034 | H | -0.300124 | 4.449936 | -2.622714 |
| C | -6.909301 | -0.191253 | 0.810395 | C | -2.043573 | 3.712675 | 0.225733 |
| C | 3.774452 | -0.729095 | 1.370555 | H | -2.897571 | 4.078419 | 0.784158 |
| C | 1.216871 | -1.001655 | 1.088124 | C | -1.298080 | 2.627545 | 0.670910 |
| C | 0.646199 | 1.064923 | 0.403686 | H | -1.563761 | 2.161296 | 1.612599 |
| C | -1.662128 | 4.335291 | -0.960511 | C | -1.246382 | -0.600803 | 0.560203 |
| C | 3.988831 | -2.534923 | -0.407011 | H | -1.897437 | 0.182781 | 0.938003 |
| C | 2.417276 | -0.273303 | 0.920034 | H | -1.408254 | -1.473965 | 1.184047 |
| C | 2.091988 | 1.028101 | 0.525909 | C | -4.108759 | -0.652340 | -0.709728 |
| C | -0.168766 | 2.183683 | -0.053397 | H | -3.999512 | 0.310996 | -1.224059 |
| C | 2.995735 | 2.159835 | 0.325164 | H | -4.082838 | -0.456440 | 0.369563 |
| C | 0.419141 | -2.353870 | 3.069580 | C | -6.650259 | -0.324075 | -0.707922 |
| H | 0.910256 | -1.637131 | 3.733725 | H | -7.568066 | -0.730878 | -1.153274 |
| H | -0.642953 | -2.101991 | 3.017204 | H | -6.495806 | 0.668359 | -1.140733 |
| H | 0.504902 | -3.348812 | 3.516319 | O | -2.957802 | -2.795824 | -0.381657 |
| C | 4.268662 | -3.909655 | -0.398697 | H | -3.906909 | -3.030140 | -0.297532 |
| H | 4.731041 | -4.357969 | 0.475838 | O | -5.655569 | -2.545147 | -0.464068 |
| C | 3.948363 | -4.695585 | -1.506193 | H | -6.057580 | -2.315720 | 0.424185 |
| H | 4.170540 | -5.758090 | -1.488357 | O | -7.306956 | 0.914596 | 1.252421 |
| C | 3.335719 | -4.122419 | -2.625540 | N | 4.375974 | -1.775586 | 0.733948 |
| H | 3.080796 | -4.735547 | -3.484122 | H | 5.225697 | -2.077828 | 1.200545 |
| C | 3.063944 | -2.751871 | -2.633742 | C | 1.096393 | -2.381733 | 1.676075 |
| H | 2.604575 | -2.291613 | -3.503151 | H | 2.130715 | -2.693067 | 1.845509 |
| C | 3.399109 | -1.952554 | -1.537433 | C | 0.474469 | -3.448720 | 0.743638 |
| H | 3.212727 | -0.886531 | -1.576653 | H | -0.597725 | -3.314107 | 0.589757 |
| C | 2.659668 | 3.450942 | 0.792460 | H | 0.968987 | -3.458168 | -0.231308 |
| H | 1.723235 | 3.610824 | 1.314216 | H | 0.624404 | -4.428748 | 1.206316 |
| C | 3.544693 | 4.514396 | 0.629841 | C | -1.550181 | -0.927290 | -0.908957 |
| H | 3.278840 | 5.495825 | 1.009391 | H | -0.797094 | -1.628479 | -1.283944 |
| C | 4.770850 | 4.317880 | -0.013977 | H | -1.481698 | -0.021257 | -1.519985 |
| H | 5.455591 | 5.149702 | -0.146668 | C | -2.934244 | -1.558994 | -1.109676 |
| C | 5.115952 | 3.043918 | -0.484017 | H | -3.027133 | -1.775205 | -2.186690 |
| H | 6.063778 | 2.887122 | -0.988938 | C | -5.478501 | -1.249984 | -1.079703 |
| C | 4.244199 | 1.974154 | -0.308148 | H | -5.496419 | -1.422797 | -2.163334 |
| H | 4.520721 | 0.996544 | -0.687678 |  |  |  |  |

**Table S4:** Coordinates of the ATVwoH anion optimized in methanol via RB3LYP/6-31+G(d,p)/IEFPCM. All values in angstrom.

| Atom | X | Y | Z | Atom | X | Y | Z |
| --- | --- | --- | --- | --- | --- | --- | --- |
| F | -2.911591 | 5.105072 | -1.340920 | C | -0.232085 | 2.648249 | -1.352553 |
| O | -6.720822 | -1.504172 | 1.466889 | H | 0.535068 | 2.273234 | -2.022359 |
| O | 4.382747 | 0.194872 | 2.261997 | C | -1.087664 | 3.660917 | -1.791896 |
| N | 0.205376 | -0.228705 | 0.736299 | H | -1.003010 | 4.082429 | -2.787733 |
| C | -6.889240 | -0.417880 | 0.809901 | C | -2.219598 | 3.614213 | 0.366233 |
| C | 3.812973 | -0.466796 | 1.380006 | H | -2.992024 | 4.006276 | 1.018960 |
| C | 1.313476 | -0.982351 | 1.077178 | C | -1.347519 | 2.608868 | 0.790049 |
| C | 0.601785 | 1.061922 | 0.395428 | H | -1.438067 | 2.226867 | 1.802274 |
| C | -2.066244 | 4.117581 | -0.918994 | C | -1.167396 | -0.728943 | 0.563444 |
| C | 4.227019 | -2.328095 | -0.324417 | H | -1.851053 | 0.032111 | 0.940451 |
| C | 2.427252 | -0.153546 | 0.950055 | H | -1.293183 | -1.607299 | 1.190842 |
| C | 1.980659 | 1.146037 | 0.535002 | C | -4.049931 | -0.813337 | -0.687171 |
| C | -0.346320 | 2.105699 | -0.060327 | H | -3.934300 | 0.168402 | -1.164223 |
| C | 2.819490 | 2.342850 | 0.302488 | H | -4.033999 | -0.656567 | 0.398674 |
| C | 0.614082 | -2.434890 | 3.049656 | C | -6.592890 | -0.481722 | -0.706110 |
| H | 1.057791 | -1.697797 | 3.726480 | H | -7.502315 | -0.862609 | -1.190284 |
| H | -0.462476 | -2.244038 | 3.003279 | H | -6.426279 | 0.529733 | -1.087672 |
| H | 0.753310 | -3.428571 | 3.489389 | O | -2.896403 | -2.960337 | -0.403998 |
| C | 4.722031 | -3.642018 | -0.320880 | H | -3.843859 | -3.204448 | -0.346895 |
| H | 5.239966 | -4.016380 | 0.557901 | O | -5.603693 | -2.712995 | -0.538912 |
| C | 4.548555 | -4.464819 | -1.435147 | H | -6.021070 | -2.520862 | 0.350468 |
| H | 4.938823 | -5.478053 | -1.415335 | O | -7.308144 | 0.664780 | 1.288689 |
| C | 3.869935 | -3.992784 | -2.562888 | N | 4.486062 | -1.527240 | 0.818037 |
| H | 3.729360 | -4.633650 | -3.427767 | H | 5.352026 | -1.732117 | 1.305552 |
| C | 3.381138 | -2.682669 | -2.567024 | C | 1.285514 | -2.382753 | 1.658040 |
| H | 2.863562 | -2.297507 | -3.440779 | H | 2.339857 | -2.623517 | 1.825152 |
| C | 3.563300 | -1.846657 | -1.462953 | C | 0.743124 | -3.483242 | 0.720946 |
| H | 3.191257 | -0.830237 | -1.490106 | H | -0.335493 | -3.411483 | 0.561870 |
| C | 2.478766 | 3.587309 | 0.864846 | H | 1.236697 | -3.447289 | -0.254633 |
| H | 1.593283 | 3.664958 | 1.487858 | H | 0.945267 | -4.464415 | 1.164425 |
| C | 3.269482 | 4.717889 | 0.645105 | C | -1.489927 | -1.073031 | -0.899286 |
| H | 2.985431 | 5.666297 | 1.092733 | H | -0.733571 | -1.770993 | -1.275210 |
| C | 4.425984 | 4.629795 | -0.136626 | H | -1.425337 | -0.168901 | -1.514360 |
| H | 5.042885 | 5.507704 | -0.305086 | C | -2.870900 | -1.706336 | -1.106308 |
| C | 4.781975 | 3.398460 | -0.696254 | H | -2.969820 | -1.903943 | -2.187097 |
| H | 5.676763 | 3.315206 | -1.306889 | C | -5.415550 | -1.393226 | -1.097006 |
| C | 3.986514 | 2.270621 | -0.480523 | H | -5.418934 | -1.521690 | -2.187140 |
| H | 4.271472 | 1.326034 | -0.934231 |  |  |  |  |

**Table S5:** Coordinates of ascorbic acid and its radical corresponding to the lowest proton affinity optimized in methanol via B3LYP/6-31+G(d,p)/IEFPCM. All values in angstrom.

|  | C<sub>6</sub>H<sub>8</sub>O<sub>6</sub> |  |  | C<sub>6</sub>H<sub>7</sub>O<sub>6</sub><sup>•-</sup> | 3-OH | position |
| --- | --- | --- | --- | --- | --- | --- |
|  | X | Y | Z | X | Y | Z |
| C | 1.732673 | -1.121442 | 0.037517 | 1.696486 | -1.056929 | 0.018792 |
| C | 1.987555 | 0.305517 | -0.092773 | 1.916435 | 0.345380 | 0.065484 |
| O | 0.428463 | -1.317156 | 0.418174 | 0.347601 | -1.332120 | 0.195843 |
| O | 2.504586 | -2.045065 | -0.157602 | 2.509137 | -1.973278 | -0.156136 |
| C | 0.850271 | 0.968796 | 0.198893 | 0.720808 | 1.015066 | 0.253760 |
| O | 3.189176 | 0.806773 | -0.477857 | 3.164716 | 0.911835 | -0.115687 |
| C | -0.204282 | -0.034166 | 0.603625 | -0.342761 | -0.077749 | 0.393805 |
| O | 0.637153 | 2.291594 | 0.184709 | 0.415105 | 2.253140 | 0.289918 |
| C | -1.491253 | 0.060295 | -0.223930 | -1.470505 | 0.092994 | -0.636161 |
| O | -3.805847 | -0.611458 | -0.442660 | -3.289351 | -0.757284 | 0.766394 |
| C | -2.575739 | -0.922087 | 0.220000 | -2.610228 | -0.910782 | -0.480920 |
| O | -1.939695 | 1.408401 | -0.028218 | -2.039638 | 1.396602 | -0.480203 |
| H | 3.798179 | 0.064143 | -0.623901 | 3.793706 | 0.185007 | -0.246780 |
| H | -0.448099 | 0.070101 | 1.668412 | -0.766450 | -0.080107 | 1.404618 |
| H | -0.330233 | 2.444840 | 0.190510 | -1.296892 | 1.999544 | -0.232308 |
| H | -1.249929 | -0.102852 | -1.283923 | -1.038960 | -0.025384 | -1.642569 |
| H | -3.799267 | -1.001417 | -1.328283 | -3.310756 | -0.775462 | -1.316932 |
| H | -2.262982 | -1.953322 | 0.027307 | -2.223541 | -1.932286 | -0.514446 |
| H | -2.768903 | -0.802305 | 1.290241 | -3.479077 | 0.189654 | 0.858817 |
| H | -2.844799 | 1.474624 | -0.372721 |  |  |  |

## Notes

### Competing Interest Statement

The authors have declared no competing interest.

